# The *Plasmodium falciparum* Drugome And Its Polypharmacological Implications

**DOI:** 10.1101/042481

**Authors:** Yinliang Zhang, Li Xie, Lei Xie, Philip E. Bourne

## Abstract

Malaria is a disease contracted by over 200 million people each year, mostly in developing countries. The primary causative agent, *Plasmodium falciparum* (*P. falciparum*) has shown increased resistance to existing drugs, hence new treatments are needed quickly. To this end we performed a high-throughput systems-level analysis, mapping existing FDA drugs with the potential for repurposing against targets from the *P. falciparum* structural proteome. The resulting *P. falciparum* drugome (*P.falciparum*-drugome) was used to prioritize potential new anti-malaria candidate targets and highlight some novel FDA approved drugs that have apparent anti-malaria effects for possible use as multi-target therapeutics.

## INTRODUCTION

The emerging field of systems pharmacology is changing the way we think about drugs. The traditional “one-drug one-target one-disease” paradigm has proven inadequate for drug discovery^1^; whereas the idea of multi-target drug interactions leading to the observed phenotype is now widely accepted^2^. Stated another way, a drug response is a consequence of complex interactions between multiple intracellular and extracellular components. Thus designing drugs assuming multiple targets is becoming the new rational approach to drug discovery, but requires a system level view of drug action. Systems pharmacology provides that view through a systematic understanding of drug action by integrating systems biology (including biological network analysis^3^), bioinformatics and cheminformatics approaches.

Simultaneously, network analysis approaches of a different kind have proven useful for organizing high-dimensional biological datasets and extracting meaningful information. Such network approaches in systems pharmacology can provide a global view of drug relationships, identify new drug targets as well as therapeutic strategies, and improve our understanding of the side effects and alternative uses of current drugs ^3-4^. Here we exploit such a network approach in determining the *P. falciparum* drugome, a structural proteome-wide drug-target interaction network that forms a basis for exploring alternative treatments for malaria.

Malaria is one of the most devastating and widespread tropical parasitic diseases and is most prevalent in developing countries. The World Health Organization has estimated that over 200 million cases of malaria occur annually ^5^; in 2010, around 655,000 deaths were reported, but this is likely a significant underestimate. Approximately 87% of these deaths were children under the age of five ^6^. To reduce the number of malaria infections and to reduce the death toll is an urgent priority.

Malaria is caused by the Plasmodium parasite, which is transmitted by a mosquito vector. In humans, the parasites multiply in the liver, and then infect red blood cells. There are five species that are known to infect humans: *P. falciparum*, *Plasmodium vivax*, *Plasmodium ovalae*, *Plasmodium malariae* and *Plasmodium knowelsi*. Among these five species, the parasite *P. falciparum* is the most dangerous, with the highest rates of complications and mortality. Currently, there are no approved vaccines available and given increasing drug resistance, finding novel anti-malarial drugs and associated targets appears the most efficient way to fight malaria^7^.

Previously Gamo et al. ^8^ screened nearly 2 million compounds from the GSK chemical library, of which more than 13 thousand were confirmed to inhibit *P. falciparum* growth. Jensen et al. ^9^ mapped the genome of *P. falciparum* to this drug-like chemical space using the chemical-protein annotation from the ChemProt database. However, establishing a new drug from such a large list of promising compounds is a lengthy, expensive and inefficient process ^10^. As the pharmacologist and Nobel laureate James Black said, “the most fruitful basis for the discovery of a new drug is to start with an old drug” ^11^. This was the approach taken by Kinnings et al. who reported a TB-drugome constructed from FDA-approved drugs and *Mycobacterium tuberculosis* (M.tb) proteins, and revealed that one-third of the FDA-approved drugs had the potential to be repositioned to treat tuberculosis and many currently unexploited M.tb receptors might be chemically druggable and could serve as novel anti-tubercular targets ^12^.

In the spirit of Kinnings et al. ^12^ the objective of the work described here is to provide a systems level view of malaria where all accessible putative target proteins are associated with all FDA-approved drugs. Hence, we built a structural proteome-wide drug-target network combining *P. falciparum* proteins with FDA-approved drugs, referred to as the *P. falciparum* drugome (*P.falciparum*-drugome). Using the *P.falciparum*-drugome we prioritized potential anti-malaria candidate targets and highlighted some novel drugs that have apparent anti-malaria effects as multi-target therapeutics. We regard the *P.falciparum*-drugome as a high-throughput approach to hypothesis generation, where the most promising theoretical drug target interactions can be tested experimentally.

## RESULTS

### A drug binding site database

A total of 308 different drugs approved for human use were identified in the RCSB Protein Data Bank (PDB) ^13^. Of these drugs, 135 were co-crystallized within a single protein structure, while the rest were co-crystallized with at least two protein structures (Figure 1), bringing the total number of drug binding sites in the PDB to 1423.

**Figure 1.**
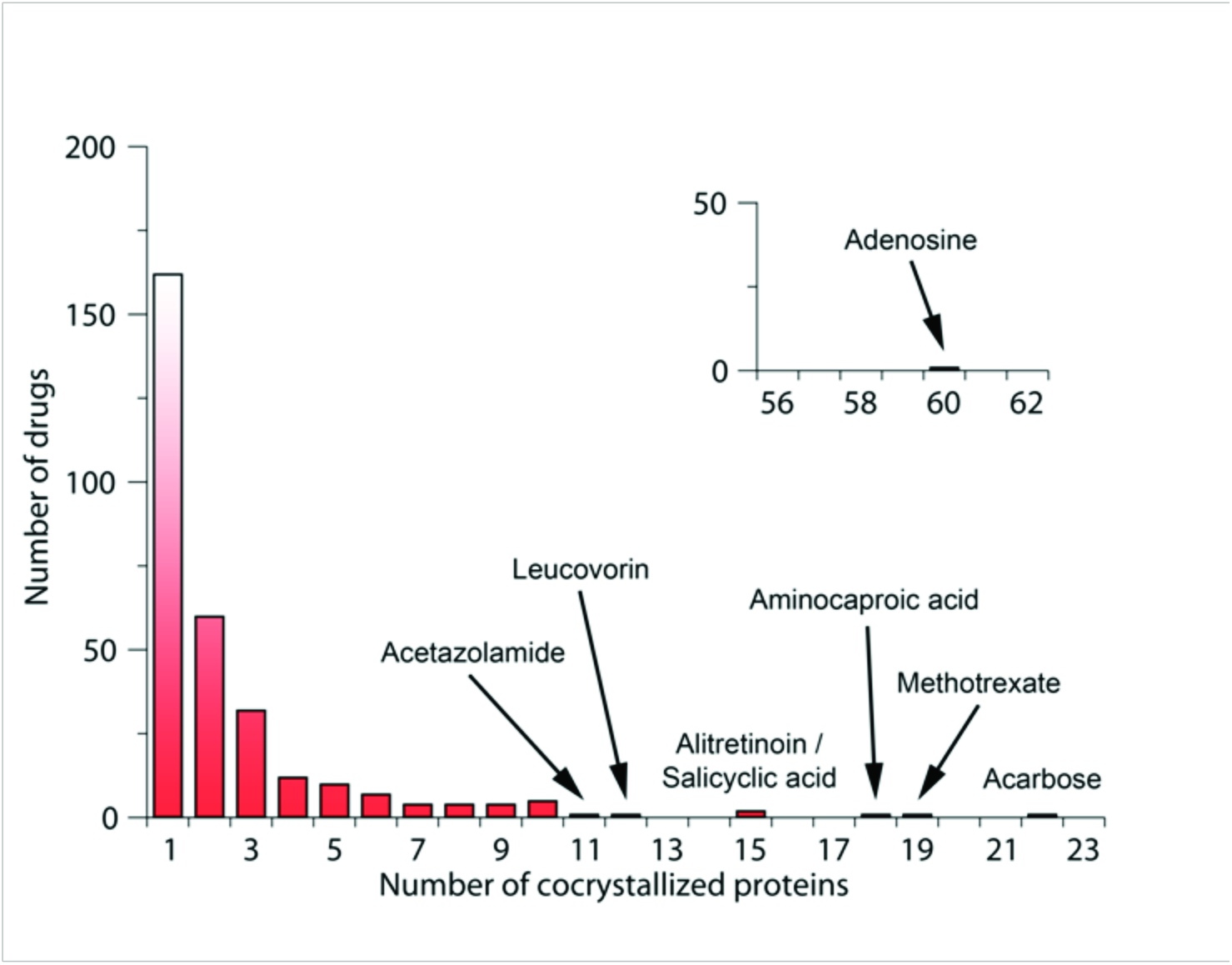
The numbers of unique proteins co-crystallized with approved drugs in the PDB.

### The *P.falciparum*-drugome: A drug-protein network

In the *P.falciparum*-drugome, 116 drugs are connected to 268 *P. falciparum* proteins (Figure 2). The *P.falciparum*-drugome is a proteome-wide drug-protein interaction network, which consists of 384 nodes and 1120 edges (Figure 2). Each node represents either an FDA-approved drug or a *P. falciparum* protein; an edge is formed when a *P. falciparum* protein has a similar binding site to the known drug target as found in the PDB (see Methods). Of 308 drugs, 116 are connected to 268 *P. falciparum* proteins that originate from a total of 1569 *P. falciparum* solved structures or models.

**Figure 2.**
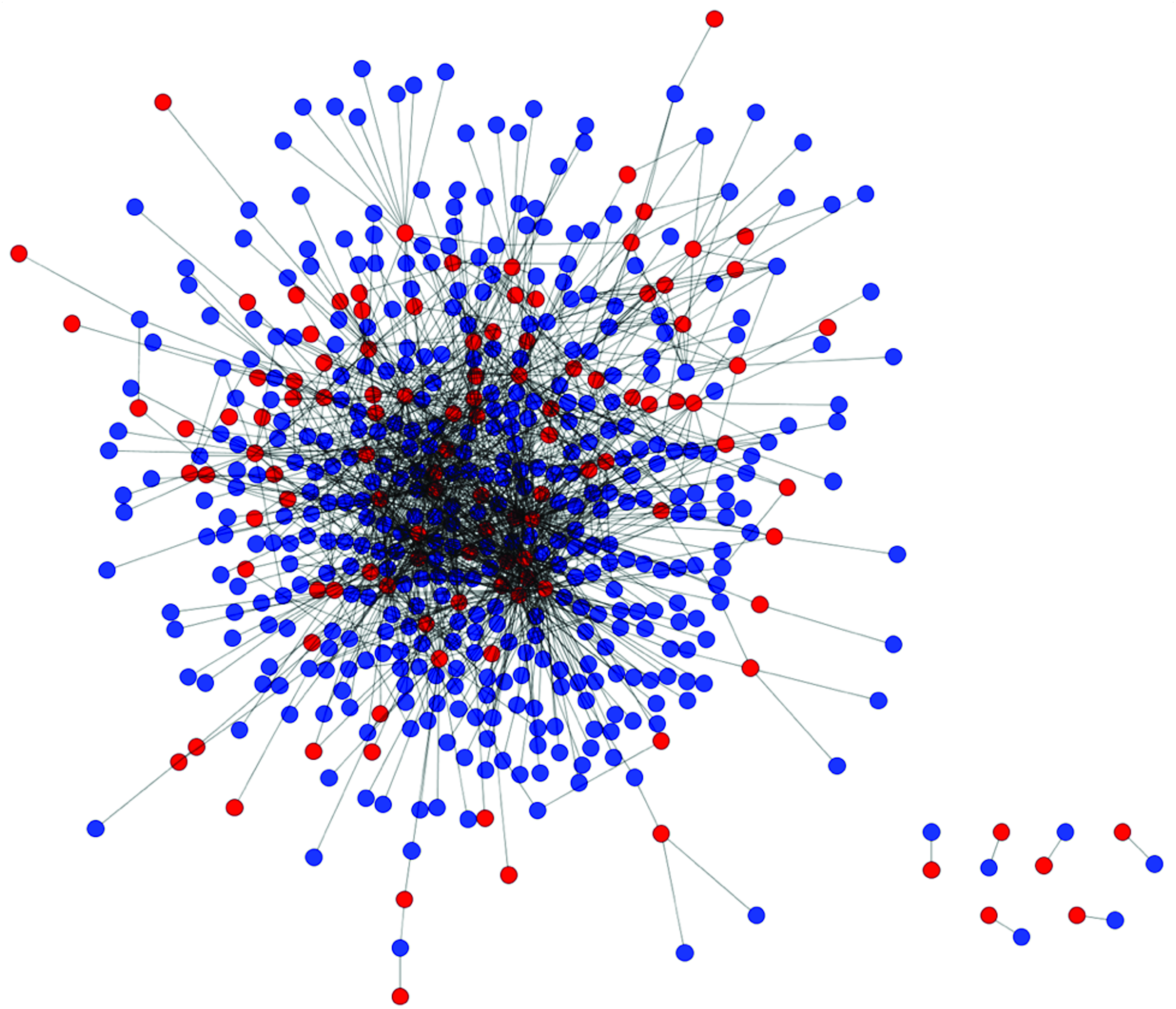
Protein-drug interaction network. The similarities between the binding sites of *P. falciparum* proteins (blue), and binding sites containing approved drugs (red) were illustrated. An SMAP p-value threshold of 1.0e-5 was used to define similarity.

We used SMAP^14^ to compare all PDB identified drug binding sites to all *P. falciparum* proteins with available structures and then subsequently excluded target-protein pairs whose global structures are significantly similar (see Methods). The average connectivity of the drugs was calculated in order to determine the p-value threshold for SMAP. As shown (Figure 3), an inflection point occurs at a p-value of 1.0e-5, that is, when the p-value is less than 1.0e-5, the drug connectivity only changes slightly, but it increases rapidly when the p-value is larger than 1.0e-5. Xie et al. previously reported that the false-positive rate is approximately 5% when the SMAP p-value is close to 1.0e-5 ^14b^. The correlation between p-value and drug degree observed here is consistent with that observed for the TB-drugome ^12^ and hence a p-value < 1.0e-5 is considered a good measure of true binding site similarity.

**Figure 3.**
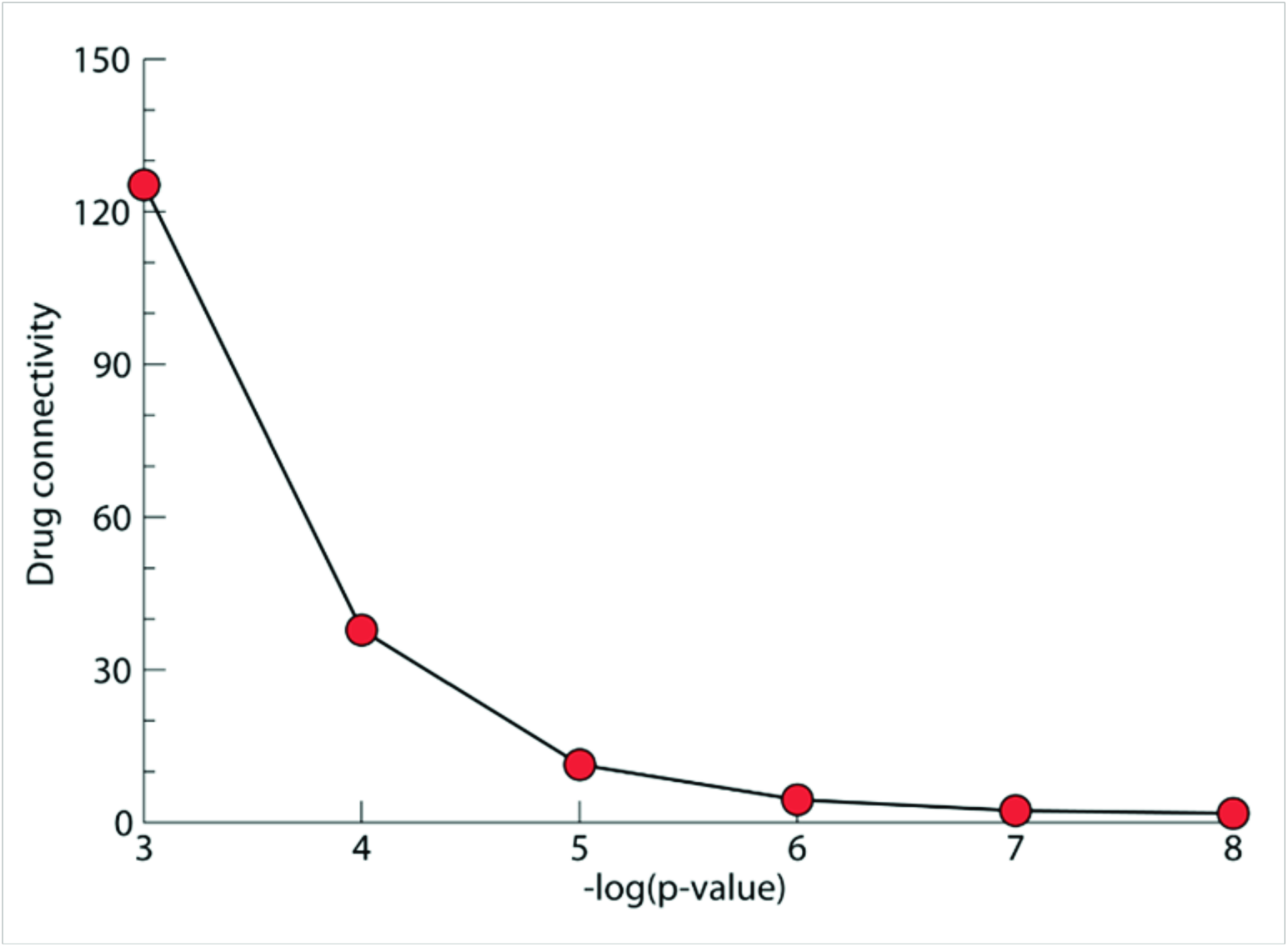
The average number of connections per drug in the P.falciparum-drugome plotted against the SMAP p-value threshold.

### The *P.falciparum*-drugome is scale-free

The distribution of target connectivity follows a power-law distribution (Figure 4), indicating that the *P.falciparum*-drugome is a scale-free network rather than random. That is, a small number of *P. falciparum* proteins are connected to a large number of drugs while the majority of *P. falciparum* proteins have few connections. The TB-drugome is also a scale-free network ^12^, suggesting that being scale-free is a common feature of these drugome networks and speaks to either a consistency in the variability of ligand binding sites across species, a limitation in the chemical space explored by existing FDA approved drugs, or both.

**Figure 4.**
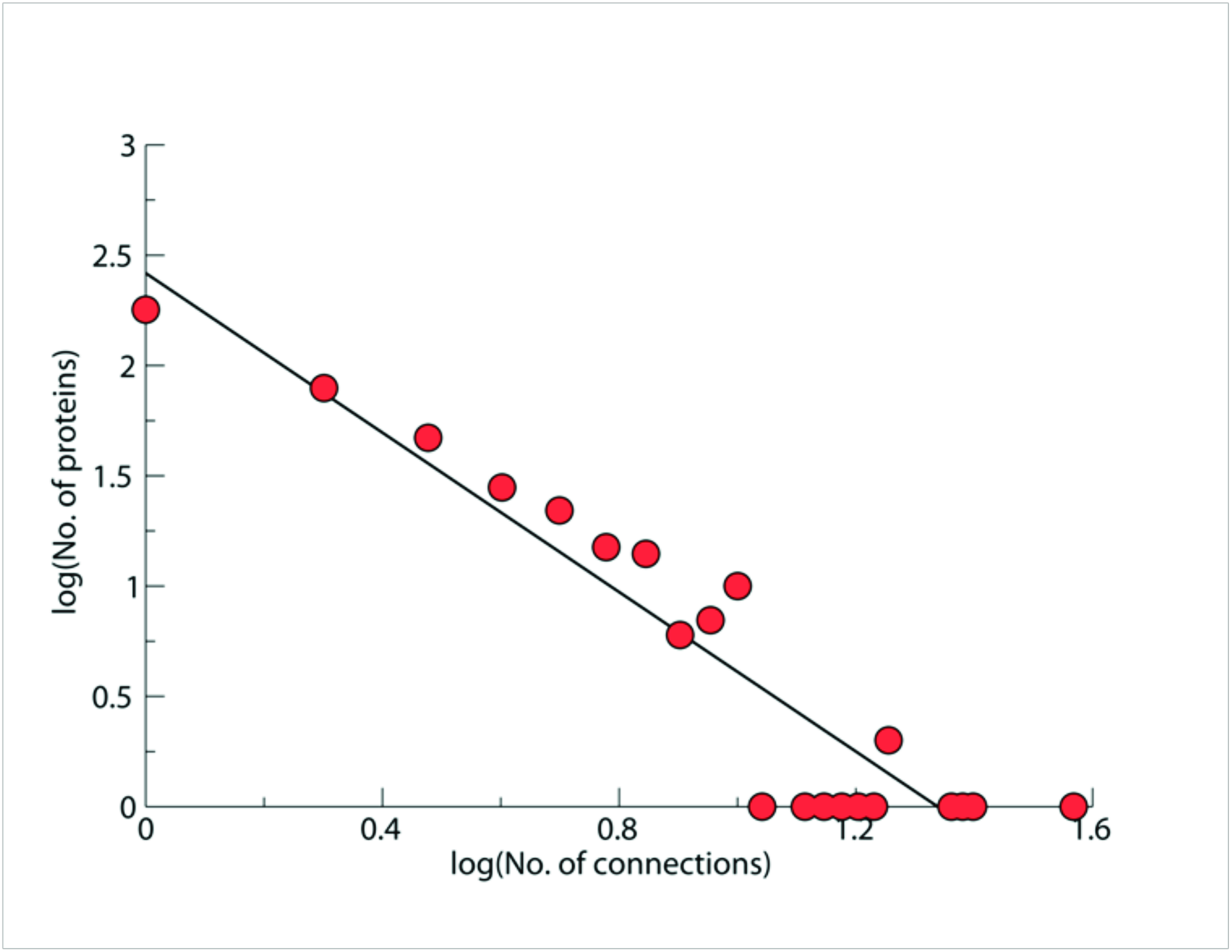
Fitting of the distribution of target connections to a power-law distribution for the *P.falciparum*-drugome. An SMAP p-value threshold of 1.0e-5 was used.

### Antimicrobial drugs are most likely to be anti-malarial drugs

Given the distribution by species of FDA approved drugs bound to their primary targets as found in the PDB and the inferred targets in the *P.falciparum*-drugome, an interesting question is whether drugs targeting a specific species are favored against *P. falciparum*. Figure 5 illustrates the affected organisms based on drug binding as found in the PDB (Figure 5A) and drugs binding to proteins in the *P.falciparum*-drugome (Figure 5B), respectively. Based on information from DrugBank, 74% of approved drugs target humans and other mammals; this number decreases to 62% in the *P.falciparum*-drugome. As expected, the distribution of drugs that primarily affected on bacteria, virus, fungi and protozoa increase from 11%, 3%, 3% and 3% to 13%, 13%, 7% and 5% respectively, indicating a higher propensity for antimicrobial drugs to bind to *P. falciparum* proteins.

**Figure 5.**
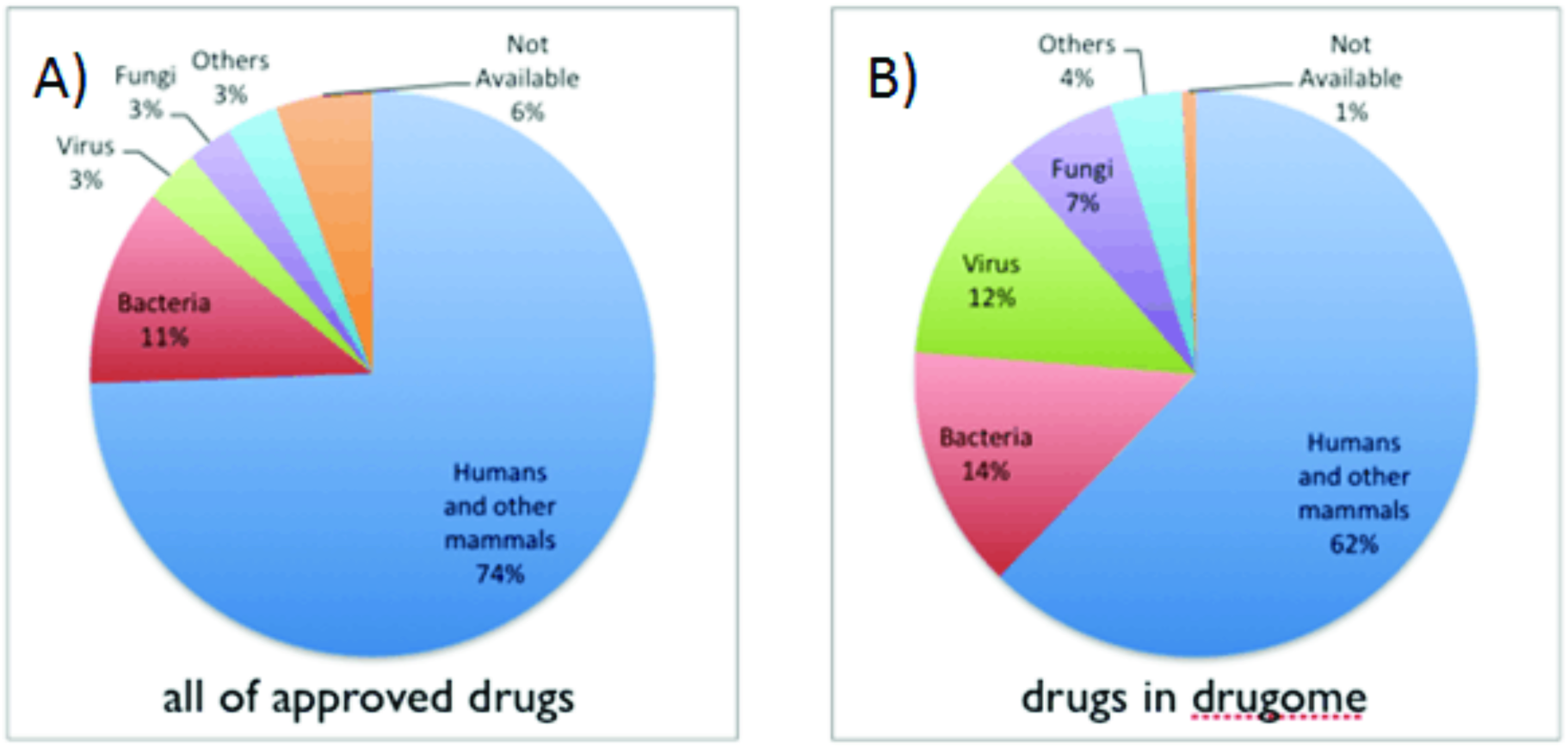
Drug distribution based on affected organisms. A) All approved FDA drugs. B) Drugs forming the P.falciparum-drugome.

Among all those antimicrobial drugs, the distribution of antiviral drugs increased most significantly, from 3% to 13% (Figure 5). Further investigation shows that 12 out of 16 antiviral drugs in *P.falciparum*-drugome are anti-HIV drugs. The association between HIV and malaria has always been an interesting issue, and many HIV protease inhibitors do have antimalarial activity^15^. Several HIV-1 protease inhibitors are proved inhibit the growth of *P. falciparum* in vitro at clinically relevant concentrations while the mechanisms are poorly understood^16^. According to *P.falciparum*-drugome, the binding sites are similar between the targets of these antiviral drugs and *P. falciparum* proteins. This may explain why most anti-HIV drugs can also inhibit malaria, yet more studies are required to verify this hypothesis.

### Highly connected drugs are candidates for multi-target therapeutics

The *P.falciparum*-drugome reveals that, of the 308 different drugs investigated, 88 drugs could potentially inhibit more than one P. falciparum protein. This is advantageous because multi-target therapeutics can be more efficacious and less vulnerable to adaptive resistance^17^. Highly connected drugs have the potential to inhibit a large number of different *P. falciparum* proteins simultaneously, a literature research of the highly connected drugs also support this hypothesis.

The top nine highly connected drugs are further revised. With 76 cross-fold connections, alitretinoin, a drug used to treat cutaneous lesions in patients with Kaposi’s sarcoma, is the most highly connected drug. Alitretinoin, also called 9-cis-retinoic acid, is a metabolite of vitamin A. It is well known that supplementation with vitamin A can significantly reduce the malaria infection and decreases the mortality of infected young children^18^. With 70 cross-fold connections, methotrexate, an anti-cancer drug, is the second most highly connected drug. Note that although serum albumin is listed as one intended target of methotrexate, this drug has not actually been crystallized with serum albumin in PDB, so this does not account for its high connectivity. Several groups have shown that methotrexate is extremely effective against P*. falciparum* in vitro ^19^. Nzila group has tested methotrexate in four murine species, however, none of them were susceptible to methotrexate. Noteworthy that they also tested the efficacy of pyrimethamine, a known antimalarial drug, in combination with folic acid in *Plasmodium berghei*, a murine plasmodium species that highly similar to human parasite *P. falciparum*, and data indicate that folic acid does not influence pyrimethamine efficacy, which suggests that the murine plasmodium, *P. berghei*, may not transport folate. Since methotrexate utilizes folate receptor/transport to gain access to cells, its lack of efficacy against the four tested murine malaria species may be the result of inefficiency of drug transport^20^. It is worth mentioning that a phase I trial in Kenyan adult healthy volunteers was carried out, and result suggests that low-dose methotrexate had an acceptable safety, but methotrexate blood levels did not reach the desirable concentration to clear malaria infection, further dose finding is necessary^21^. With 63 different cross-fold connections, levothyroxine, a drug used to treat hypothyroidism, is the third most highly connected drug. Further investigation revealed that it was the structure of levothyroxine bound in the binding site of serum albumin that was determined to be significantly similar to 44 of the 63 different *P. falciparum* binding sites. As a non-specific binder of steroid hormones and a transport protein for various fatty acids, serum albumin is known to be a highly promiscuous protein^22^. To our knowledge, there is no report of levothyroxine having any effect on malaria. Besides, the forth highly connected drug, raloxifene, a drug used to prevent and treat osteoporosis and breast cancer in women, also hasn’t been reported affecting malaria. The fifth highly connected drug, estradiol, is a sex hormone. In the past few years, evidence has accumulated that many hormones, especially the sex steroids, can influence the immune system and thus susceptibility for diseases caused by protozoan parasites^23^. An *in vivo* experiment shows that estradiol can decreases parasitemia, but increases the incidence of cerebral malaria and the mortality in *P. berghei* infected mice^24^. Although the complete mechanism has not yet been unraveled, there is still an evidence that estradiol has effect on *P. falciparum* proteins.

It is worth to notice that drugs with an above-average number of connections are either antibacterial drugs or human affecting drugs (Figure 6). Although the whole distribution increased slightly, part of the antibacterial drugs are connected to a larger number of different *P. falciparum* proteins than antiviral drugs, making these certain antibacterial drugs the most likely candidates against malaria assuming the goal is to impact as many targets as possible in the battle to overcome resistance through mutation.

**Figure 6.**
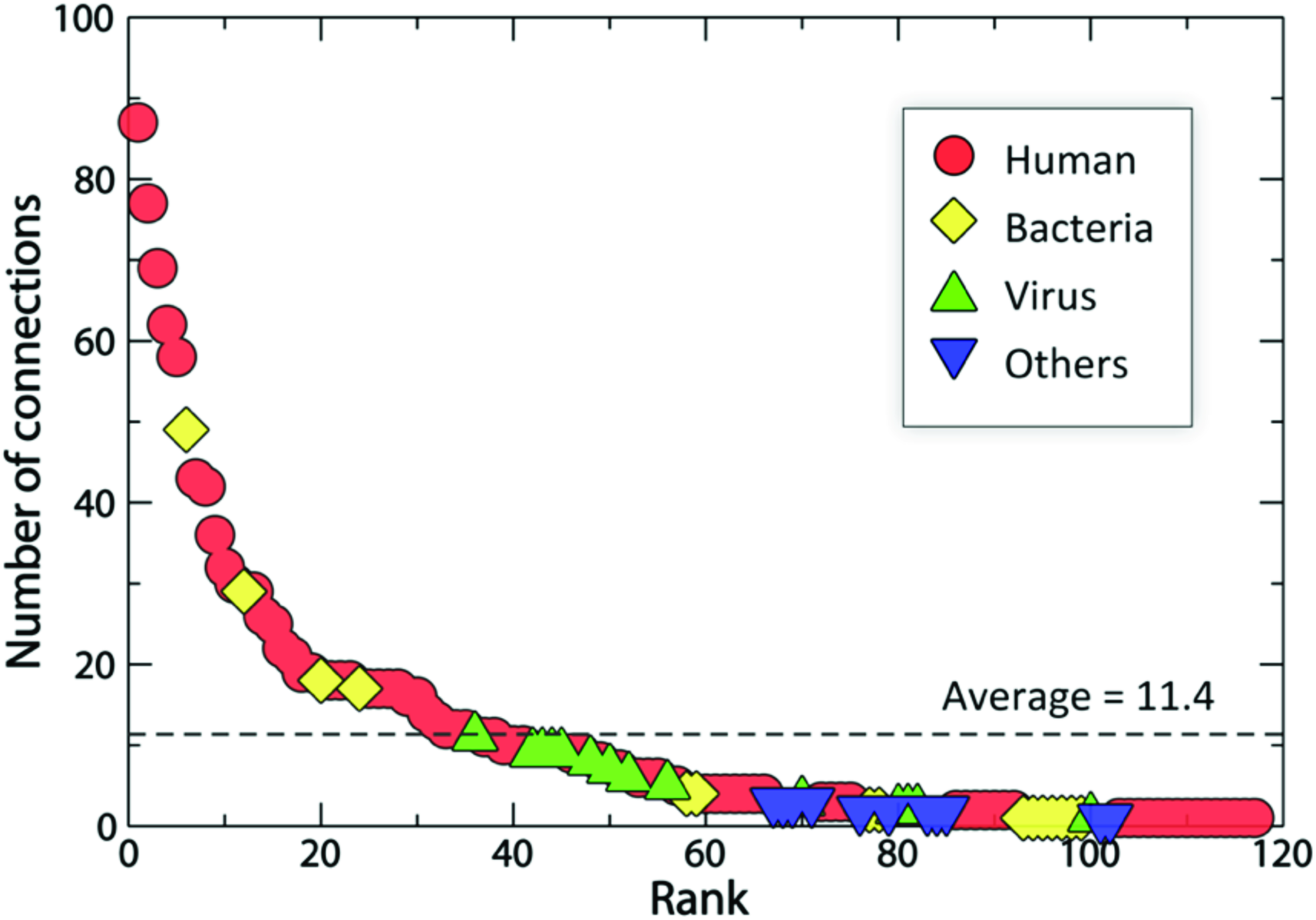
A ranking of FDA approved drugs according to the number of targets to which the drug is assumed to bind.

The sixth highly connected drug, fusidic acid, is a bacteriostatic antibiotic. Many studies suggest that fusidic acid has potential activity against malaria^25^, Johnson et al. shows that fusidic acid inhibited the growth of *P. falciparum* erythrocytic stages with immediate death effect^26^. Other antibacterial drugs whose connections are above average are rifampin, trimethoprim and tetracycline. The antimalarial activity of rifampin was confirmed with *P. falciparum* in vitro^27^, and a combination with other 2 drugs was verified safe and efficacious for treating malaria in murine model^28^. Trimethoprim is a pyrimidine inhibitor of dihydrofolate reductase, and World Health Organization (WHO) guidelines for the treatment of malaria recommend a combination of trimethoprim and sulfamethoxazole as prophylaxis for HIV-infected patients to protect against malaria^29^. Tetracycline is a broad spectrum antibiotic, and has also been recommended by WHO as second-line antimalarial treatment when combined with artesunate or quinie^29^.

To further investigate the effects of gene encoding P. falciparum proteins in metabolism, we here use a genome-scale flux-balanced metabolic network model^30^. We revised top 9 highly connected drugs and the *P. falciparum* proteins they connected to which are predicted essential for growth in the unconstrained metabolic network of *P. falciparum*. All of these 9 drugs are predicted bind at least one essential *P. falciparum* protein, indicating the highly connected drugs have high potential to be antimalarial drugs.

The concept of “synthetic lethality”-some genes may not essential on their own, but are lethal if deleted simultaneously – sheds new light on drug development^31^. The most interesting thing is that, in the *P. falciparum* metabolic network, they predict 16 gene pairs that have synthetic lethality, and one of them occurs in alitretinoin, levothyroxine and mifepristone. Another pair of the synthetic lethal genes is connected to different drugs, making it possible to use these drugs as combination therapy.

According to these experimental supports, it is highly believed that with the help of our discovery the representative drugs listed in our *P.falciparum*-drugome are worth exploring and testing thoroughly.

## DISCUSSION

Existing drug-target networks are typically constructed from either annotated drug-target pairs or transference of chemical properties. Here we present our second (the first was the TB-drugome) high throughput structural-based proteome-wide drug-target network, that of P. falciparum – the *P.falciparum*-drugome. By using Food and Drug Administration (FDA) approved drugs we present the raw data for possible repurposing strategies, since these drugs are assumed safe. By presenting the drug target interaction network based on binding site similarity across much of the *P. falciparum* proteome we provide raw data on possible new targets for these drugs where the hubs represent the best opportunities to combat resistance. Experimental support exists for some of our theoretical findings and a variety of putative new targets and repurposing opportunities are presented in the supplemental materials. Simultaneously we are beginning to examine the TB-and *P. falciparum* drugomes against the emergent human drugome. This will be the subject of a future publication. Likewise, we are working on a workflow software system that will enable others to simply compute drugomes or update existing drugomes as new drugs and putative targets become available.

It is worth mentioning that Yuan et al used a high-throughput chemical screening approach, and identified 32 highly active compounds^32^. Medicines for Malaria Ventrue (MMV) also provides a resource named Malaria Box, which contains 200 drug-like compounds and 200 probe-like compounds, and they are selected by *in vitro* screening against P. falciparum 3D7 from 20,000 compounds in GlaxoSmithKline Tres Cantos Antimalarial Set (TCAMS), Novartis-GNF dataset and St. Jude Children’s Research Hospital’s dataset ^33^. The compounds they provide are different from the drugs in our drugome, mainly because most of the compounds are not approved drugs or have no co-crystallized structure as yet. While chemical high throughput screening will find compounds with direct antimalarial effect, some of the most promising candidates for combined therapy may be missed. For example, the highly connected drug in our drugome, rifampin, which is proven safe and efficacious for treating malaria in a murine model when combined with isoniazide and ethambutol, but not alone ^28^, does not exist in either Yuan’s work or Malaria Box.

Notwithstanding there are some unavoidable limitations to our approach. First, although the genome of *P. falciparum* is sequenced ^34^, structural coverage is only 2.1% from experimental structure and 12.5% when homology models are included. Models may be of limited accuracy, being modeled from single chains, missing many protein-drug interactions that occur at the interfaces between tertiary units of the protein and only observed if the quaternary structure is modeled. Even more limiting is the bias in the PDB from which experimental structures are drawn and models inferred. Many drug targets, notably membrane proteins, are underrepresented in the PDB. While the number of FDA approved drugs rises slowly the structural coverage of a variety of proteomes, including human is increasing more rapidly and will only see this type of high throughput technique improve, especially when used with emergent methods from systems pharmacology.

## MATERIALS AND METHODS

### Structural coverage of the *P. falciparum* proteome

There are 5491 protein-encoding genes in the *P. falciparum* proteome ^34^, 118 of which have solved structures available in the RCSB Protein Data Bank ^13^ (Oct 18, 2011). A single protein sequence may have several associated structures co-crystallized with different ligands, or containing different point mutations. All are important in comparing binding sites, which brings the total number of known *P. falciparum* protein structures to 333 (Oct 18, 2011).

To maximize coverage of the P. falciparum proteome we also include homology models deposited in ModBase ^35^, a database of annotated comparative protein structure models and associated resources. This database contains over 20 thousands homology models for the entire P. falciparum proteome since ModBase has several models for each protein sequence; a model is considered reliable if its Model score (GA341) is greater than 0.7 and its ModPipe Protein Quality Score (MPQS) is greater than 1.1 (http://modbase.compbio.ucsf.edu/modbase-cgi/display.cgi?type=help&server=modbase). Employing these thresholds, there are 1236 reliable homology models (Oct 18, 2011), and the total structural coverage of the P. falciparum proteome is 12.5%. As indicated in the Discussion, only a single chain of each homology model is available, rather than the entire biological unit.

### Identification of FDA-approved drug binding sites

The DrugBank database ^36^ combines detailed drug and target information where known (http://www.drugbank.ca/). Drugs approved by the U.S. Food and Drug Administration (FDA) are taken from DrugBank and all the nutraceuticals removed. We use InChI keys to map these FDA-approved drugs to ligands in the PDB to find the associated co-crystalized structures. After removing non-protein structures, such as DNA, RNA and associated complexes, there are 308 FDA-approved drugs with 1423 drug-protein binding sites (Dec 6, 2011).

### Comparison of ligand binding sites using SMAP

Xie et al. developed SMAP ^14^ for the comparison of potential protein three-dimensional ligand binding site motifs in a manner that is independent of the sequence order. Proteins are represented by their shapes, as defined by using only Cα atoms. Then ligand binding sites of the two proteins are aligned and assigned a p-value measuring their similarity based on a unified statistical model ^14b^.

We compare 1423 drug binding sites against 1569 *P. falciparum* proteins, including both experimental structures and homology models. The entire structure of each *P. falciparum* protein is used to avoid missing alternative binding sites on any given *P. falciparum* protein. A *p*-value is reported to measure the similarity of the binding sites for each pair.

### Comparison of global protein structures using FATCAT

FATCAT (Flexible structure AlignmenT by Chaining Aligned fragment pairs allowing Twists) ^37^ is an approach for flexible protein structure comparison. The Java version of FATCAT was downloaded from the RCSB PDB website and run locally (http://pdbx.org/jfatcatserver/download.jsp).

In order to detect only remote similarities between the *P. falciparum* proteins and drug binding site proteins, that is, those which cannot be identified through global structure comparison, we exclude those pairs identified by FATCAT where the p-value was less than 0.5.

### Visualization of the protein-drug interaction network

We use yEd from yWorks to visualize the protein-drug interaction network (http://pdbx.org/jfatcatserver/download.jsp).

In order to detect only remote similarities between the *P. falciparum* proteins and drug binding site proteins, that is, those which cannot be identified through global structure comparison, we exclude those pairs identified by FATCAT where the p-value was less than 0.5.

### Visualization of the protein-drug interaction network

We use yEd from yWorks to visualize the protein-drug interaction network (http://www.yworks.com/en/products_yed_about.html). A node represents a *P. falciparum* structure or a drug ligand. If a *P. falciparum* structure and a drug binding site have similar binding sites, and their global structures are dissimilar (according to the criteria outlined above), we place an edge between the *P. falciparum* structure and the associated drug.

### GO term enrichment analysis

To evaluate the functional relationships between target proteins of each highly connected drug, we use DAVID (Database for Annotation, Visualization and Integrated Discovery) ^38^ to perform a GO term enrichment, since DAVID provides a comprehensive set of functional annotation tools for investigators to understand biological meaning behind large list of genes. DAVID uses a modified Fisher Exact p-value to measure the enrichment of each GO term. The smaller a *p*-value is, the more enriched this GO term is.

We use the UniProt accessions of each target protein and associated drug binding sites to do the GO term enrichment. For each highly connected drug, we establish the GO term enrichment of the associated *P. falciparum* structures. In general terms, if these targets are in diverse pathways, this drug may hold promise for a multi-target treatment regime, with particular emphasis on avoiding resistance. Alternatively, if these targets are enriched in a specific pathway, this means the drug have the potential to treat the specific disease without side effects.

## Supporting Information

**Table S1**. Information about the approved drug binding sites used in the P.falciparum-drugome. This file contains information about the 308 approved drugs that were identified in the PDB. For each drug, its name, PDB ligand code, isomeric SMILES string and known targets are listed, and the PDB codes of the protein structures with which it has been crystallized are given.

**Table S2**. Cross-fold drug-target pairs in the P.falciparum-drugome (for solved P. falciparum structures only). This file contains a list of the cross-fold drug-target pairs with a SMAP P-value <1.0e-5, for solved P. falciparum structures only. For each pair, information about the drug and target structures is given, as well as the corresponding SMAP P-value (indicating the significance of the binding site similarity) and AutoDock Vina score (from docking the drug into the predicted binding site in the P. falciparum protein).

**Table S3**. Cross-fold drug-target pairs in the P.falciparum-drugome (for P. falciparum homology models only). This file contains a list of the cross-fold drug-target pairs with a SMAP P-value <1.0e-5, for homology models of P. falciparum proteins only. For each pair, information about the drug and target structures is given, as well as the corresponding SMAP P-value (indicating the significance of the binding site similarity) and AutoDock Vina score (from docking the drug into the predicted binding site in the M.tb protein).

**Table S4**. Information about the solved P. falciparum structures used in the P.falciparum-drugome. This file contains information about the P. falciparum proteins with solved structure(s) in the RCSB PDB that were used in the P.falciparum-drugome. For each protein, the gene name, protein name and corresponding PDB codes are given.

**Table S5**. Information about the P. falciparum homology models used in the P.falciparum-drugome. This file contains information about the reliable homology models of P. falciparum proteins from ModBase that were used in P.falciparum-drugome. For each homology model, the ModBase model code is given, as well as the gene name and description of the P. falciparum protein.

## ACKNOWLEDGMENT

We thank Dr. Haiyan Liu for his helpful discussions and suggestions. Financial supports from China Scholarship Council are gratefully acknowledged.

